# Uncovering convergence and divergence between autism and schizophrenia using genomic tools and patients’ neurons

**DOI:** 10.1101/2023.08.11.552921

**Authors:** Eva Romanovsky, Ashwani Choudhary, David Peles, Ahmad Abu Akel, Shani Stern

## Abstract

Autism spectrum disorders (ASDs) are highly heritable and result in abnormal repetitive behaviors and impairment in communication and cognitive skills. Previous studies have focused on the genetic correlation between ASDs and other neuropsychiatric disorders, but an in-depth understanding of the correlation to other disorders is required. We conducted an extensive meta-analysis of common variants identified in ASDs by genome-wide association studies (GWAS) and compared it to the consensus genes and single nucleotide polymorphisms (SNPs) of Schizophrenia (SCZ). We found approximately 75% of the SNPs that are associated with ASD are also associated with SCZ. We have also probed cellular phenotypes reported in ASD neurons compared to SCZ neurons from induced pluripotent stem cell (iPSC) models. Interestingly, Autism neurons start with an early maturation and schizophrenia neurons start with a late maturation, but both end up with deficits in synaptic activity when compared to control neurons as they mature.

## INTRODUCTION

The nature of the relationship between autism spectrum disorder (ASD) and schizophrenia (SCZ) has been the subject of intense debate. ASDs are a complex and heterogeneous group of neurodevelopmental disorders that affect approximately 1% of children globally and vary in degree of severity^1^. According to the DSM-5 criteria, abnormal social interaction, repetitive behavior, and delayed language and cognitive skills are the main characteristic symptoms used to diagnose ASD at an early age^2^. There are several genetic and non-genetic risk factors associated with ASD. Recent studies suggest that ASD is one of the most highly heritable neuropsychiatric disorders (83%)^3^. Large-scale genomic studies (e.g., SPARK) have led to the implications of hundreds of genes that might play an important role in the manifestation of . SCZ is associated with the presence of core symptoms that have been classified along negative (e.g., blunted affect, asociality) and positive (e.g., delusions and hallucinations) dimensions, as well as cognitive disorganization, and its prevalence has been estimated at about 0.35% worldwide^5^. SCZ similarly to ASD has high heritability estimates (∼80%) with clear difference in the age of onset (manifestation of SCZ is generally in adulthood)^6^.

While diagnostically independent, clinical reports indicates that ASD co-occurs with SCZ and vice-versa at rates higher than in the general population^7^, as well as with other developmental and neurological disorders including, among others, intellectual disability, attention-deficit hyperactivity disorder (ADHD), anxiety, depression, and epilepsy^8^. Moreover, multidimensional evidence from phenotypic, behavioral, neuroimaging, environmental and genetic substrates point to considerable overlap between ASD and SCZ^7,9^. A meta-analysis by a cross-disorder group of the psychiatric genomics consortium studying eight psychiatric disorders involving subjects of European ancestry discovered the genetic correlation between the disorders in GWAS data but also classified them into three different groups^10^. ASD and SCZ were classified into different groups of "early onset neurodevelopmental disorders" and "mood and psychotic disorders" respectively^10^. Moreover, increased parental age and maternal immune activation are the environmental risk factors that are common to both disorders^1^ Besides the evidence of genetic and environmental risk factor correlations, clinical symptoms based on DSM-5 also appear to overlap, especially the impairments in social interaction in ASD and the negative group of symptoms in SCZ like social withdrawal and reduced communication^7^. The meta-analysis of neuroimaging studies has also suggested alteration of common brain regions and functional networks among neuropsychiatric disorders, including SCZ and ASD^11,12^

Given the observed overlap of certain features of SCZ and ASD, it is pertinent to dissect the neurobiological pathophysiology and disease phenotypes and seek the commonalities between the disorders. Rare structural variants like deletion of chromosome 22q11.2 and 15q13.3 or duplication of chromosome 16p11.2 are associated with both SCZ and ASD and also with other developmental disorders. But the majority of rare variants (or copy number variations) stretch across large genomic regions and hence the position of mutation at the genic or allelic level is difficult to infer. For instance, exonic deletions of NRXN1 are a known monogenic rare variation that increases the risk for ASD, SCZ, and other neurodevelopmental disorders but the position of reported exonic mutations vary among different disorders^13^. A recent study investigated the pleiotropy of rare variant genes in SCZ and neurodevelopmental disorders^13^.But more understanding of the shared genetic and pathophysiological processes is required especially for the GWAS-discovered common variants.

Over the last decade, induced pluripotent stem cells (iPSC) based models of SCZ and ASD^14–22^ have provided important clues about the pathophysiological processes affected in neural cells, especially since the animal models do not fully recapitulate the complex genetics and clinical features of neuropsychiatric disorders^23^. We recently performed a quantitative meta-analytical study of the progress done in the field of SCZ genetics and the findings of SCZ phenotypes in iPSC-derived neural cells^6^. Some prior studies have focused on the shared rare variants and their pleiotropic effects in the development of SCZ and ASD^24–26^.

Here in this study, we have analyzed genome-wide association studies (GWAS) performed on ASD individuals to find the frequently reported consensus genetic variants. We have also compared the consensus gene variants of ASD (single nucleotide polymorphism, SNPs) with the frequently reported genetic variants in SCZ^6^ to determine the shared and specific SNPs (gene variant) affected and prevalent in both disorders to have an in-depth understanding of the genomic correlation between ASD and SCZ. In addition, to investigate the functional role of ASD GWAS genes, we intersected them with Developmental Brain Disorder Database (DBD) genes and traced them to determine the brain regions with the greatest percentage of ASD common variants. Furthermore, we have performed a comprehensive analysis of approximately 50 iPSC-based studies of ASD to uncover the highly common disease phenotypes and have attempted to correlate them with the commonly reported ASD genes by performing gene set enrichment analysis. Hence, in this study, using a quantitative meta-analytical approach to genomics and an iPSC-based disease modeling, we have probed both the converging and distinct features between ASD and SCZ.

## METHODS

### Genetic Variant Analysis and Standardization of Gene Nomenclature of Autism Spectrum Disorder (ASD) and Schizophrenia (SCZ) Genes

The genetic variant analysis and the standardization of gene nomenclature were performed according to Choudhary et al., 2022^6^. Trait statistics were retrieved from the NHGRI-EBI Genome-Wide Association Study (GWAS) Catalog^27^ for the study of ASD (EFO_0003756) and SCZ (EFO_0004609). A total of 17 publications with 305 genes related to ASD and 86 publications with 1119 genes related to SCZ were reported. The *SNP_GENE_IDS* column was utilized to identify the Ensembl IDs of genes that contain ASD-related reference SNPs (rs, SNP-risk allele) within their sequence. When several Ensembl IDs were attributed to one gene in a single publication, they were unified and the gene symbol was counted once in each publication to reduce noise from the data because Ensembl IDs of the same gene symbol had overlapping loci in most cases. Furthermore, care was taken to ensure that each publication was counted only once since some publications had multiple entries and thus the same Ensembl IDs. For each gene variant, the frequency was calculated by counting how often it was reported in a publication with a *p*-value of <0.01. The most frequently annotated genes for ASD are those counted in more than three publications (out of n=23) and for SCZ those counted in more than five publications (out of n=105).

The full dataset from the Developmental Brain Disorder Gene Database (DBD)^28^ was downloaded to extract a list of genes associated with ASD. According to DBD, 672 genes have been linked to autism. The Ensembl Data (Biomart, 2022-08-25)^29^ was accessed to acquire a list of Ensembl IDs and their corresponding gene name. The chosen dataset was *Human Genes (GRCh38.p13)* from *Ensembl Genes 107*and the attributes selected were *Gene Stable ID* and *Gene Name*. Additionally, the database of Human Gene Nomenclature Committee (HGNC) gene symbolsfromEMBL-EBI Public FTP[http://ftp.ebi.ac.uk/pub/databases/genenames/hgnc/tsv/hgnc_complete_set.txt] (2022-08- 25) was downloaded which contained approved gene symbols, previous gene symbols, alias gene symbols and the related Ensembl ID for each gene. The gene nomenclature was standardized for GWAS and DBD gene symbols using HGNC as described in Choudhary et al.2022^6^. The method involves searching for the Ensembl ID in both the Biomart results and the HGNC database. If the Ensembl ID is found in the HGNC database, the matching approved symbol from the HGNC database is used. If the Ensembl ID is not found in the HGNC database, the Biomart symbol is compared to the list of approved HGNC symbols. If the symbol is an approved HGNC symbol, it is used. If the symbol is not an approved symbol, it is searched in the list of previous gene symbols in the HGNC database. If the symbol matches a previous symbol of a gene, the currently approved HGNC symbol is used. This method facilitates the matching of HGNC-approved symbols to all Ensembl identifiers with no potential for confusion.

For ASD genes of GWAS, the identifier *LINC02163* was replaced by the approved symbol *NIHCOLE*. Similarly, for ASD genes of DBD, the identifier *C12ORF57* was not located in either the HGNC nor the Biomart symbols, and the identifier *KIAA0100* was substituted by the approved symbol *BLTP2*. The gene symbols of SCZ were already standardized by Choudhary et al., 2022^6^.

### Data analyses

All statistical analyses were performed by using R (version 4.1.2 (2021-11-01))^30^and MATLAB (version 9.12.0.2009381 (R2022a) Update 4, 2022-07-7)^31^.

### Investigating the Commonalities between ASD and SCZ Genes; Single Nucleotide Variants (SNPs)

To find common genes, the intersection of ASD genes from GWAS and DBD was formed first (n=30 genes), followed by most frequent ASD genes and SCZ genes from GWAS (n=17 genes). Overlapping genes were represented as word clouds using MATLAB. Genes were color-coded so that their overlap with the corresponding gene list could be seen. The frequency of genes associated with ASD that appeared more than three times in publications was visualized using MATLAB.

Subsequently, the commonalities between ASD and SCZ were further investigated. For this purpose, the most frequent genes of ASD (≥3 publications) and SCZ (≥5 publications) were overlapped to create a common gene list. The SNPs of the common genes as reference SNP (rs) were taken from the GWAS catalogs of ASD and SCZ (*SNPS* column). The common genes were matched with the SNPs and merged into one heatmap. ASD (red) and SCZ (green) were marked when the SNP was detected. The *oncoPrint()* function from the R package *ComplexHeatmap*^32^ was used to generate the heatmap. In addition, all chromosomes were represented as a circular layout to mark the genomic position of detected SNPs. The human reference genome assembly GRCh38 was used. The *RCircos* R package^33^ was employed for this purpose.The maturation and neuron development scheme in iPSC models of ASD and SCZ has been simplified and created with Biorender.com The scheme is referred to^15,18,34,35^.

### Analysis of RNA Expression Levels of ASD Genes Across Brain Regions

RNA consensus tissue gene data (*rna_tissue_consensus.tsv*) was downloaded from The Human Protein Atlas (HPA, proteinatlas.org)^36^. This data set provided a summary of normalized expression levels for genes across 54 different tissues, based on transcriptomics data from HPA and Genotype-Tissue Expression (GTEx). The data included an Ensembl gene identifier (*Gene*), analyzed sample (*Tissue*) and normalized expression value (*nTPM*) for each gene. It was based on the HPA version 22.0 and Ensembl version 103.38.

First, the data set was subset by the overlapping ASD genes of GWAS and DBD. The average gene expression was calculated for each tissue, filtered by brain regions including the amygdala, basal ganglia, cerebellum, cerebral cortex, hippocampal formation (hippocampus), hypothalamus, medulla oblongata (medulla), midbrain, pons, spinal cord, and thalamus. BrainNet ^37^ and <u>Biorender.com</u> were utilized to assign color-coding based on the ranking of average expression level, with red signifying a high expression level and violet representing a low expression level. The illustration is not a precise representation of the anatomical structure of the brain but rather a schematic one. The colored areas were used to demonstrate the different brain regions.

### The Genetic Pathways of ASD Explored Using Gene Set Enrichment Analysis

A gene set enrichment analysis was performed on the most commonly observed ASD genes and those ASD genes which overlapped in both GWAS and DBD to ascertain any biological connections between them and the genetic pathways correlated to ASD. The CytoScape STRING app^38^ was used to create a gene set network and to perform the functional enrichment analysis. The terms of 15 categories, including Gene Ontology Cellular Component (GO) and Monarch Phenotypes (EFO, HP), were analyzed. The results of the analysis were considered significant when the false discovery rate (FDR) adjusted *p*-value was less than 0.05 using the Benjamini-Hochberg procedure. Significant terms associated with ASD or SCZ were highlighted in the gene set network.

### Disease Phenotypes in ASD-Induced Pluripotent Stem Cells (iPSC) Through Data Visualization

A total of 51 ASD-iPSC-related publications were evaluated (Supplementary Table 1). Listed studies by Chiffre et al.^39^ were also included. The overview of numerous collected data was divided into several pie charts. Subsequently, the individual phenotypes and their morphological and functional changes during the differentiation of ASD-iPSCs were compiled as a heatmap. The regulation or effect strength has been color-coded. The green color represents low regulation (↓) and the red color represents high regulation (↑) of reported disease phenotypes. The color bar represents the number of ASD-iPSC-based models in which the corresponding regulation was observed. The R programming language was used to create the pie charts and heatmap, and the R packages *ggplot2*^40^ and *ggpubr*^41^ were used for this purpose.

## RESULTS

### Linking the genome-wide associated genes of ASD and SCZ

Using the GWAS Catalog^27^, we searched the database for "Autism Spectrum Disorders" and identified the highly associated genes when collecting the data from 17 studies.We included the studies from diverse/different ethnic groups/ancestry as plotted in Supplementary Fig. 1A. A substantial number of studies were from European ancestry followed by African and East Asian ancestry.

Fig. 1A shows the genes of the significantly associated variants with ASD that appeared in publications with a *p*-value of <0.01; The genes are presented in a word cloud. The frequency of genes (the number of publications in which the gene was reported) was denoted by the size of the font. We then checked which of the genes that showed associations with ASD are also associated with SCZ using the GWAS Catalog as a reference database as analyzed in our previous study^6^. These genes have been marked in green in Fig. 1A.

**Figure 1:**
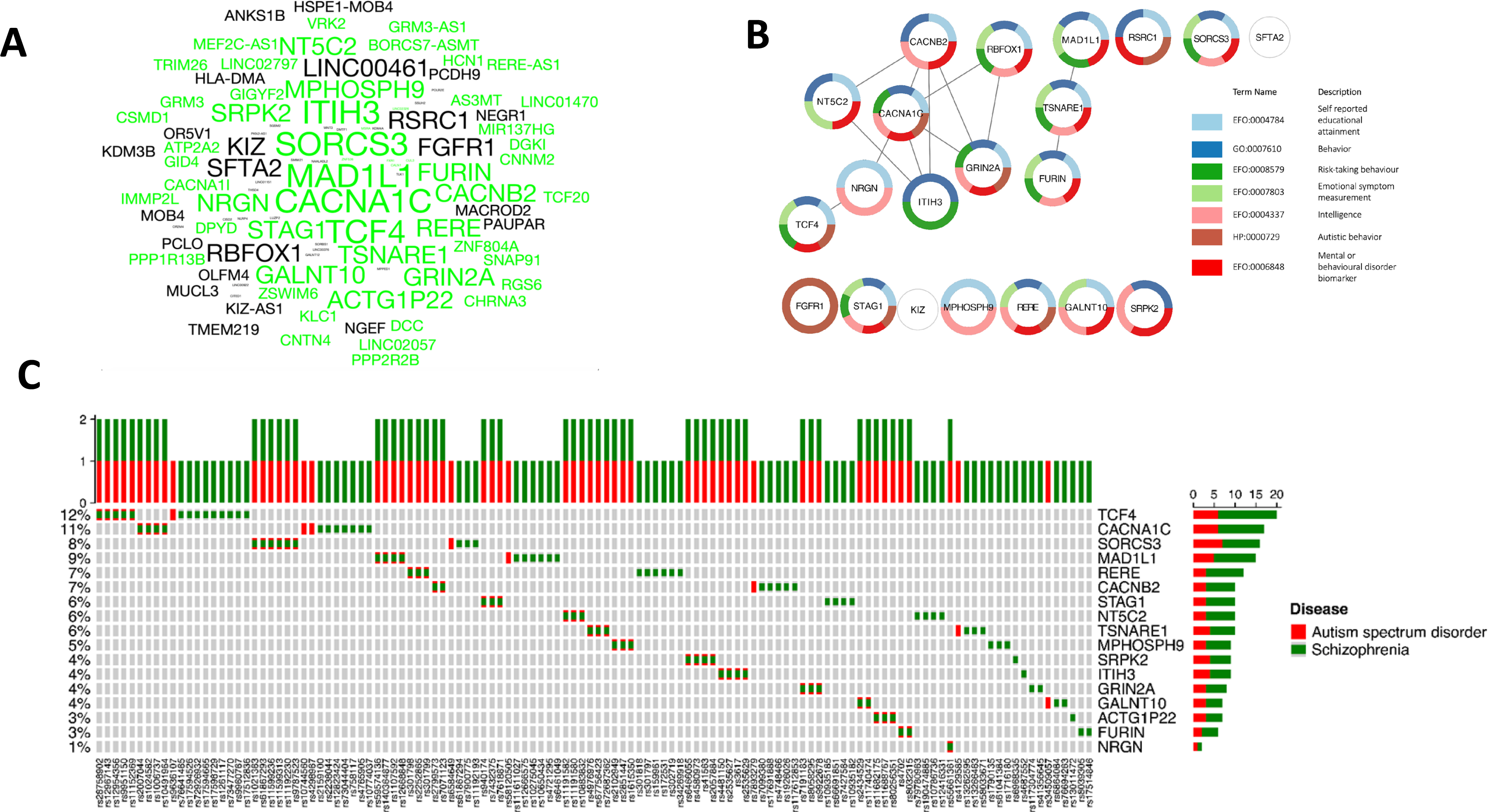
ASD and SCZ GWAS genes. (A) The word cloud of GWAS-reported genes associated with ASD. GWAS-reported genes associated with SCZ are highlighted in green. The font size indicates the frequency of gene mentions in publications, with a larger font size indicating a higher number of reported occurrences.The most frequently reported genes associated with ASD have been plotted in Supplementary Fig. 2A. (B) Gene network of the most reported genes in GWAS associated with ASD. Functional analysis revealed significant pathways indicating autistic behavior. Connected genes imply interactions. (C) Color map for shared SNPs between ASD and SCZ in GWAS. The intersection of the most reported genes in ASD and SCZ and their 122 SNPs has been visualized.

Out of all 305 reported ASD genes in GWAS, 239 genes (78%) were also associated with SCZ, and 78 were associated with the 105 (74%) most frequently reported SCZ genes in GWAS (Fig. 1A). Remarkably, 74% of the most associated genes in ASD are also associated with SCZ (Supplementary Fig 2A). The eight most frequently reported genes (MAD1L1, CACNA1C, SORCS3, TCF4, ITIH3, STAG1, TSNARE1, MPHOSPH9) were reported in more than three publications and are also associated with SCZ (Supplementary Fig 2A). Some of the most associated genes are linked to neuron function. These include subunits of calcium channels (CACNA1C, CACNB2, CACNA1l), TSNARE1 (a part of the SNARE complex), GRIN2A (a subunit of the NMDA receptor), MAD1L1 (Mitotic Arrest Deficient 1 Like 1), CACNA1I, SORCS3, SPRK2, ITIH3, and more.

We then examined the functional linkage of the 23 genes with the highest prevalence in ASD GWAS and visualized a gene network as shown in Fig. 1B. For this purpose, weused the STRING functional enrichment analysis.16 genes exhibited significant enrichment for "Self-reported educational attainment" (*p* = 4.3e-17), and 15 genes demonstrated analogous enrichment for "Behavior" (*p* = 3.9e-16). CACNA1C was found to be directly connected to other five genes (CACNB2, GRIN2A, NRGN, RBFOX1,ITIH3). Other genes like TCF4 and NT5C2 also formed a part of the network. Most of these genes are known to be associated with calcium-mediated activity of neural cells^42^. The common significant terms ascribed to these genes were “Intelligence”,“Mental or behavioral disorder biomarker”,“Risk-taking behavior” etc. The genes MAD1L1, TSNARE1, and FURIN formed another distinct network as they are part of secretory pathways including synaptic vesicle release and fusion. Some of the terms associated with these genes were “Risk-taking behavior”, "Intelligence”, and “Mental or behavioral disorder biomarker”.

### Specific SNPs that are common between SCZ and ASD

Subsequently, we aimed to investigate whether the type of mutation determines the specific brain disorder (ASD vs. SCZ) in the 17 associated genes that were common between ASD and SCZ (Fig. 1C). For example, for the TCF4 gene (a transcription factor), nine SNPS are associated with SCZ only, one with ASD only, and five SNPs are common. Similarly, for the CACNA1C gene, seven SNPs are associated with SCZ only, two with ASD only, and four with both ASD and SCZ. Overall, more SNPs are associated with SCZ and common SNPs. There are also a few SNPs associated exclusively with ASD, possibly attributed to the comparatively fewer ASD GWAS conducted compared to SCZ. As mentioned earlier, ASD patients are known to have an increased risk of SCZ^9^ and Fig. 1C provides additional support for this assertion, as the majority of SNPs linked to ASD are also associated with SCZ.

We also mapped the regions in the different chromosomes that are associated with ASD-specific genetic variants (Suppl Fig. 3A) and both with ASD and SCZ (Suppl Fig. 3B). Chromosome 3 (10 SNPs), 7(9 SNPs), and 10 (9 SNPs) had the highest number of specific SNPs reported for ASD. Chromosome 7 and 10(9 SNPs each) had the highest number of common SNPs shared in ASD and SCZ. Overall 63 SNPs from 17 genes were common between ASD and SCZ out of total 82 SNPs from 23 genes reported in ASD (Suppl Fig. 3).

### Common genes between ASD GWAS and Developmental Brain Disorder Database (DBD)

Moreover, we sought to investigate the overlap between our catalog of genes identified in ASD GWAS and those present in the DBD (Fig. 2A & Supplementary Fig. 2B).The DBD includes information obtained through exome and genome sequencing, chromosome microarray analyses, and copy number variation studies. It covers six different developmental brain disorders, including ASD and SCZ, and provides comprehensive phenotypic details that aid in the detection of pathogenic loss of function variants.We identified 30 genes that were shared between ASD GWAS genes and DBD genes. Notably, the top reported ASD GWAS genes CACNA1C, TCF4, STAG1, RBFOX1, and RERE were among this shared gene list.

**Figure 2:**
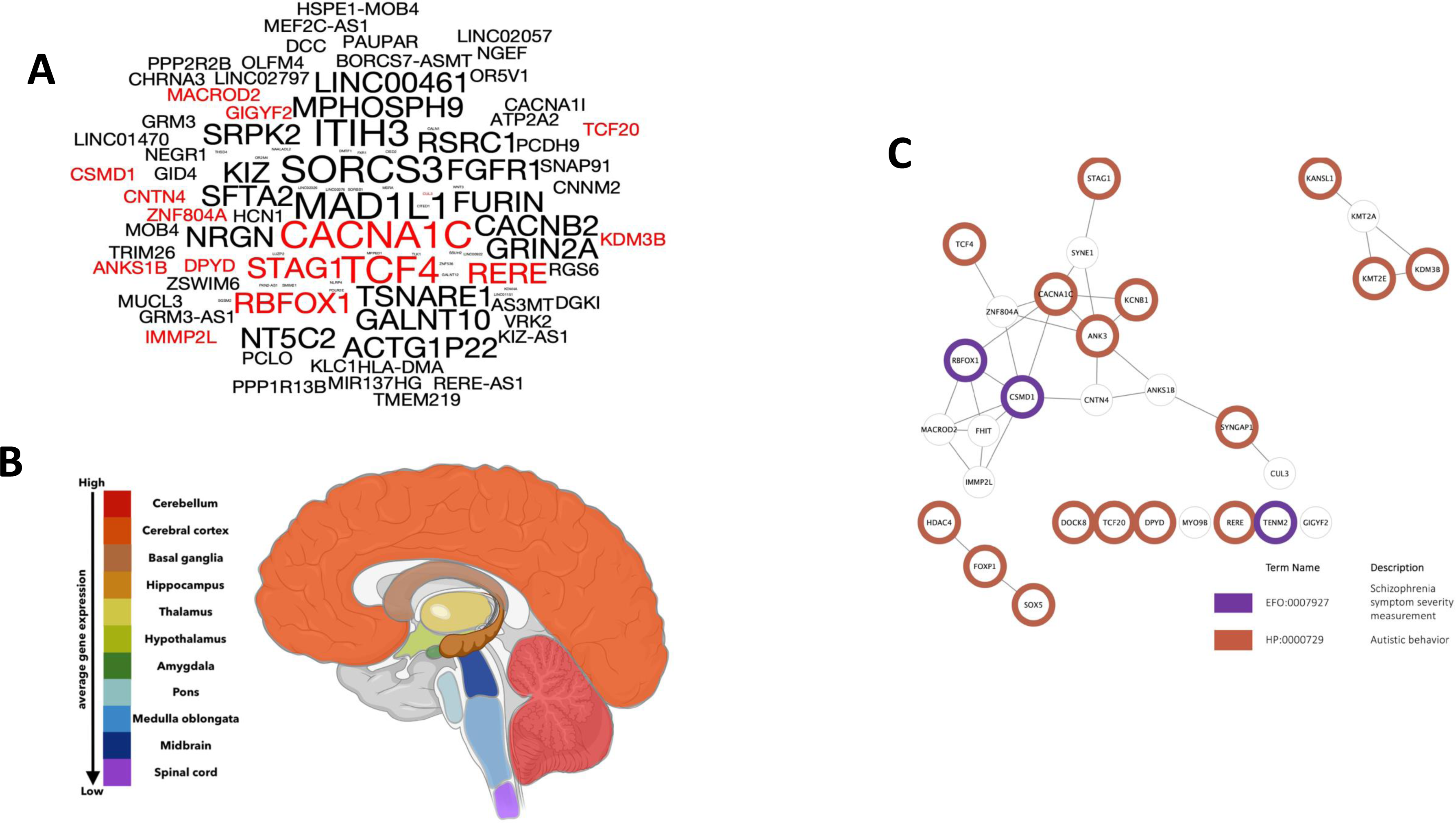
ASD GWAS and DBD genes. (A) The word cloud of GWAS reported genes associated with ASD.Genes cataloged in the DBD are highlighted in red. The font size indicates the frequency of gene mentions in publications, with a larger font size indicating a higher number of reported occurrences.The highly frequently reported genes have been plotted in Supplementary Fig. 2B. (B) RNA expression levels of ASD genes across brain regions. Overlapping genes of GWAS and DBD were used for creating the gene list (n=30) and the average gene expression level of each brain region was calculated.The “hot” colors represent higher expression levels of the specific brain region and “cold” colors lower expression levels. The graphical image was created with BrainNet ^37^ and <u>Biorender.com</u>. (C) Gene network of the common gene list of GWAS and DBD. Color-labeled genes are significantly enriched in either category of autism (*p*=6.82e-19) or schizophrenia (*p*=2.9e-04).

We next used the RNA consensus tissue gene data to determine the brain regions that may be severely impacted by ASD from human protein atlas (HPA). With the overlapping gene list of genes shared between ASD GWAS and DBD, the average gene expression was computed for each tissue.The findings were represented using color-coded visualization, demonstrating the distinct brain regions influenced by ASD (Fig. 2B). The cerebellum exhibited the highest expression level, while the spinal cord displayed the lowest expression level, with a 37% decrease compared to the cerebellum.

Furthermore, we analyzed which of the genes were associated with autistic vs. schizophrenic behaviors as illustrated in Fig. 2C. 16 genes exhibited significant enrichment for "Autistic behavior" (*p* = 9.7e-16). In contrast, a more limited enrichment was found for "Schizophrenia symptom severity measurement" (*p* = 0.0003), with only three genes displaying significant associations. Thus, it appears that the resulting gene list is more likely to be associated with autistic behavior. In addition, network analysis revealed the presence of three distinct gene clusters. In particular, two smaller clusters, consisting of three and four genes, respectively, showed significant associations with autistic behavior. Conversely, the largest cluster, comprising 16 genes, showed pronounced associations with ASD and SCZ.

### iPSC models and neuronal transcriptome in ASD

We further performed a meta-analysis of previously published work using iPSC models (see methods) (Suppl. Table 1) for ASD, as performed previously for SCZ^6^. Initially, we segmented the data to see the different ASD types (Fig. 3A) . The majority of the data is classified as “ASD” (29.4%), while a large portion of the data is Rett Syndrome (27.5%), and the next largest group is fragile X (FXS 11.8%). Approximately 39.2% of the iPSC models had isogenic lines for controls (Fig. 3B). The cell type distribution from which the reprogramming was performed is shown in Fig. 3C. The vast majority of the studies started by reprogramming fibroblasts (52.9%). The reprogramming was usually performed with a retrovirus (50.9 %), or a lentivirus (35.3%),or a Sendai virus (21.6%). In the context of ASD, most research is performed on neurons or neural progenitor cells (Fig. 3E). It is interesting to note that more than 25% of the ASD patients in the studies also have epilepsy (Fig. 3F). The distribution into neuronal types of the analyzed studies is shown in Fig. 3G.

**Figure 3:**
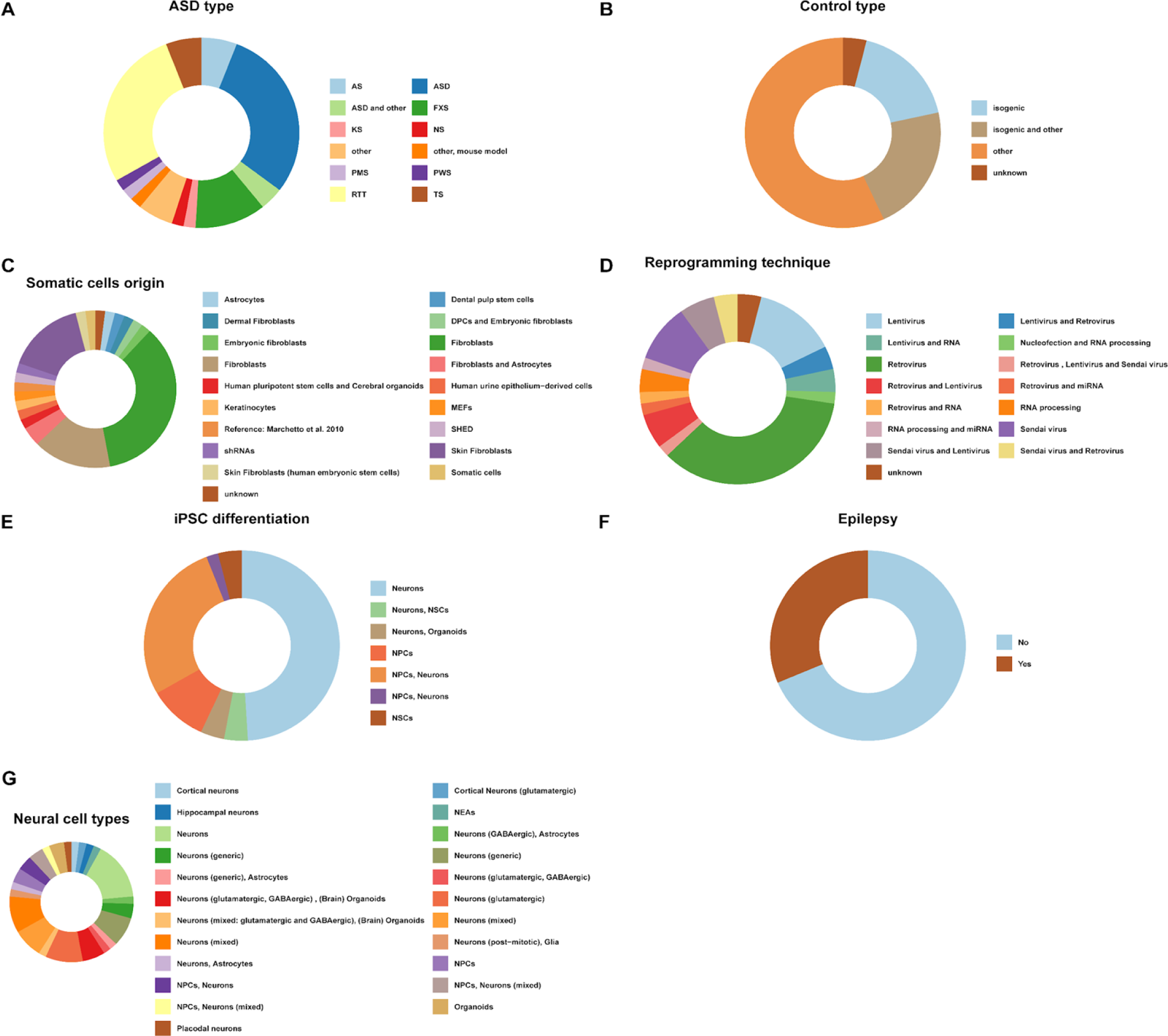
Metanalysis of iPSC models of ASD. (A-G) Donut plots of the summary statistics of the collected 51 publications of iPSC models. Seven categories *(ASD, Control, Somatic cell origin, reprogramming technique, iPSC differentiation, Epilepsy, and Neural cell types)* describe the frequency of specific patients and procedural methods in iPSC models.

Fig. 4A provides a summary of the phenotypes that were found using the iPSC models. The studies report changes such as differentiation rate, neurites’ length, and differential gene expression. Out of 51 publications, 19.6% reported low synaptic and network activity in neurons derived through NPC from ASD patients. In addition, 11.8% reported shorter dendrites and neurites and 6% reported increasing associations with mitochondrial function in the neurons derived from the ASD patients. However, the direction is reversed when specifically examining cortical neurons. Five out of seven publications reported increasing synaptic and network activity and likewise, other publications could detect increasing spike frequency and increased activity in calcium imaging^18,19,34,35^. Differential gene expression analyses showed a trend toward increased regulation of genes^18,35^.

**Figure 4:**
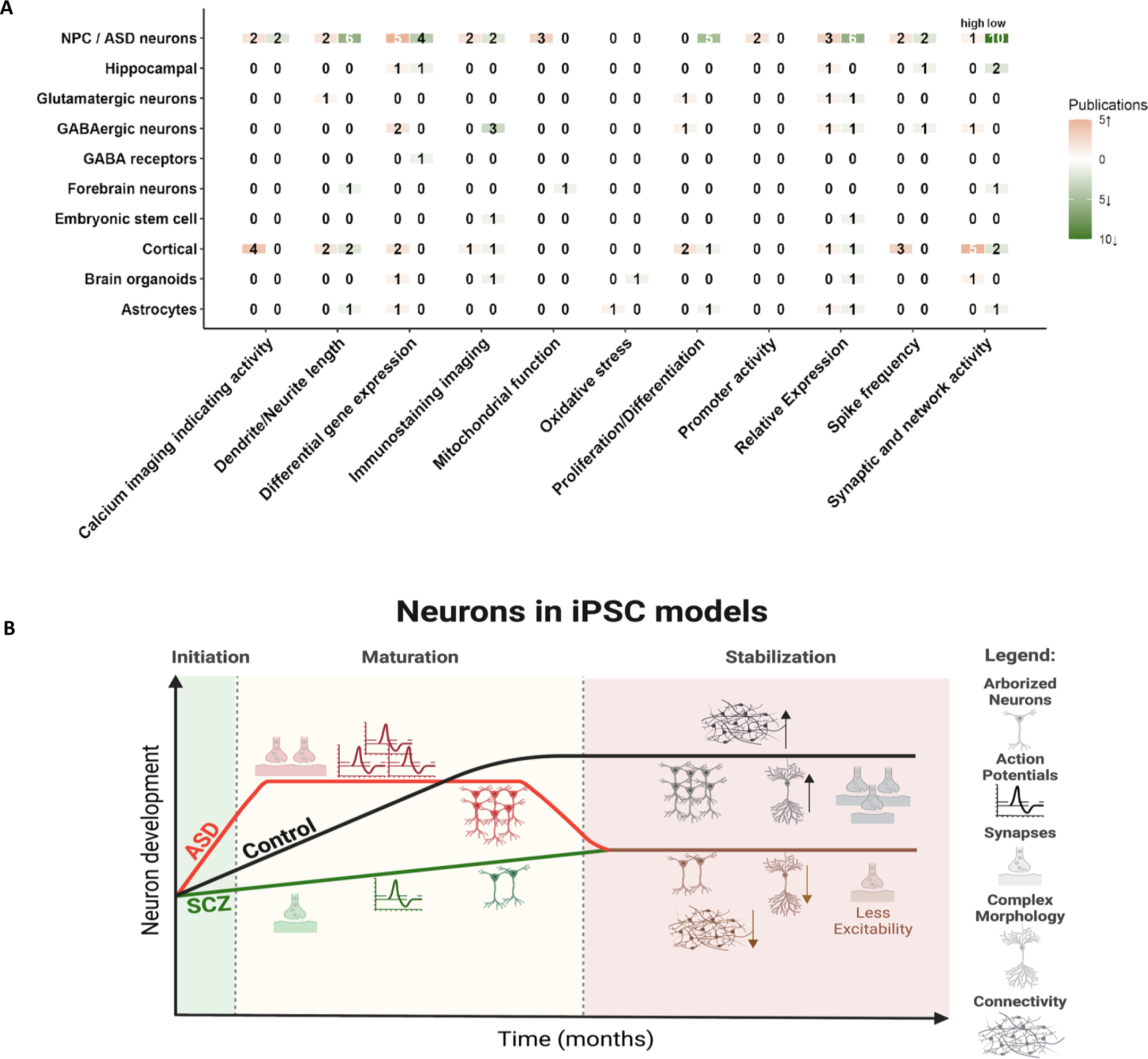
Summary of iPSC studies on ASD. (A) Heatmap of the number of publications reported activities of specific phenotypes in iPSCs models. If the number of publications is higher, the assigned color becomes darker. With reported high regulation (↑), the color becomes redder, and with low regulation, it becomes green (↓). The color white with the number “0” implies no indication. (B) Schematic of neuron progression over months in reported iPSC models^18,19,34,35^.Time was divided into three phases (initiation, maturation, stabilization). As the neurons develop, the curve also increases. The control curve describes the normal course of neuronal development and serves as a comparison for the ASD and SCZ curves. The arrows (↑↓) next to the figures describe whether this development is increased or decreased. The graphical image was created with <u>Biorender.com</u>

Fig. 4B summarizes the functional phenotypes that are usually reported. The neurons derived from ASD patients usually start with an expedited maturation^19,34^ compared to the control neurons. This includes increased sodium and potassium currents, hyperexcitability, more arborized neurites, and even more synaptic connections initially. However, as the neurons mature, they lose their synaptic connections, have reduced currents, are less excitable, and are less arborized compared to the control neurons derived from individuals without the disorder^19,35^. Interestingly, the neurons derived from the SCZ patients have a different trajectory but in the later time points, they end up with a similar phenotype^18^. The SCZ neurons start less arborized, are less excitable, have decreased sodium and potassium currents, and have less synaptic activity compared to the control neurons. When the SCZ patient-iPSC-derived neurons develop, they always lag behind the control neurons and when they are mature^18^, many of their functional and morphological phenotypes are similar to the neurons derived from the ASD patients (Fig. 4B).

## DISCUSSION

Until now, it has been difficult to determine by GWAS, the potential causal genetic variants of either ASD or SCZ. Moreover, most of the previous studies on genetic correlations of the two disorders have been focused on studies based on European ancestry. Hence, to gain better insights, we performed this study with the extensive analysis of the GWAS data on ASD individuals from diverse ancestry/ethnicity and then compared them to GWAS of SCZ individuals (also from diverse ethnicity) to find the genes and the SNPs that are common or specific to both disorders.

We found approximately (¾) fraction of the highly reported GWAS genes to be common between ASD and SCZ. Our further analysis of the SNPs of the highly reported genes revealed that most of the SNPs that are associated with ASD are also associated with SCZ. This observation may explain why ASD patients have an increased risk of SCZ ^9^.

To understand the functional role of GWAS-identified genes, we also searched for the overlap of these genes with DBD genes which has an exhaustive list of potential causal genes from six disorders including Intellectual disability, ASD, ADHD, bipolar disorder, SCZ, and epilepsy along with phenotypic information. These overlapping critical genes were expressed in different brain regions but a high percentage of them were expressed in the cerebellum, cortex, basal ganglia, and hippocampus. The cerebellum and basal ganglia are known to coordinate motor-related functions. The hippocampus and cortex are important for cognition, memory, and learning as well as language and social communication. Hence from this analysis, we can infer that variation and mutations resulting in functional dysregulation of these genes could affect these brain areas and circuits resulting in ASD symptoms.

iPSC-based models offer an alternative to understanding the effect of genomic mutations and associations on neural cells^6,16,21,43–48^. The majority of studies investigated NPCs and neurons derived from ASD patient-specific iPSC *in vitro* and only a few havestudied the role of glial cells from ASD patients using iPSC technology. The most prominent change that was common across many patient iPSC-derived neuron studies was the alteration in the synaptic activity. It may seem confounding that some of the studies reported an increase while some reported a decrease in synaptic activity. These differences could in part be explained due to the different types of neurons studied (cortical, hippocampal, gabaergic, mixed neuron cultures). Other critical reasons for the variability reported could be the timepoint of differentiation studied. Recently, Brant et al., 2021^19^ and Hussein et al., 2023^34^ have shown how neurons derived from ASD patients start with a higher network connectivity compared to a control network derived from individuals who do not have ASD, but end up when they are mature with a less connected network. From the iPSCs-based studies of ASD and SCZ, we see a different trend of phenotypes emerging for these two disorders^6,18^. The neurons from ASD patients mature earlier with increased arborizations^35^ and exhibit electrophysiological properties earlier than controls in earlier timepoints studied however gradually they show deteriorating electrophysiological properties in later timepoints. In the case of SCZ, the neurons are morphologically less arborized as well as exhibit inferior electrophysiological properties throughout the entire differentiation time compared to controls^18^ (Fig. 4B). This means that originally the two disorders start with opposite phenotypes, but eventually, the neurons derived from these two disorders have similar phenotypes when they are mature.

In conclusion, our analysis of common genomic variants and the neuronal pathophysiology of ASD in comparison to SCZ hints at the presence of an initial diverging neuronal pathophysiology that upon maturation converges into a similar pathophysiology.

## LIMITATIONS

The limitations of this study is the difference between the scale of GWAS between ASD and SCZ both in the number of studies (17 ASD studies vs. 86 SCZ studies) and also the smaller sample size recruitment in case of ASD.

## ACKNOWLEDGMENTS

The Zuckerman STEM leadership program and Israel Science Foundation grants - 1994/21 and 3252 /21 for Dr. Shani Stern.

## AUTHOR CONTRIBUTION

SS- Supervision, Conceptualization, Methodology, Data Analysis, Drafting and editing – original manuscript. AAA- Conceptualization, Editing –original manuscript, Data Analysis. ER and AC-Methodology, Data Analysis, drafting the original manuscript. All authors reviewed the manuscript. ER and AC contributed equally to the work.

