## Supplementary Figures for "Uncovering convergence and divergence between autism and schizophrenia using genomic tools and patients’ neurons"

### Supplementary Figure 1

Pie chart distribution of GWAS performed from different ethnic groups across different GWAS publications. The studies table was exported from GWAS with the key word “autism spectrum disorder” (.csv format). Overall, 45 study accessions were reported.

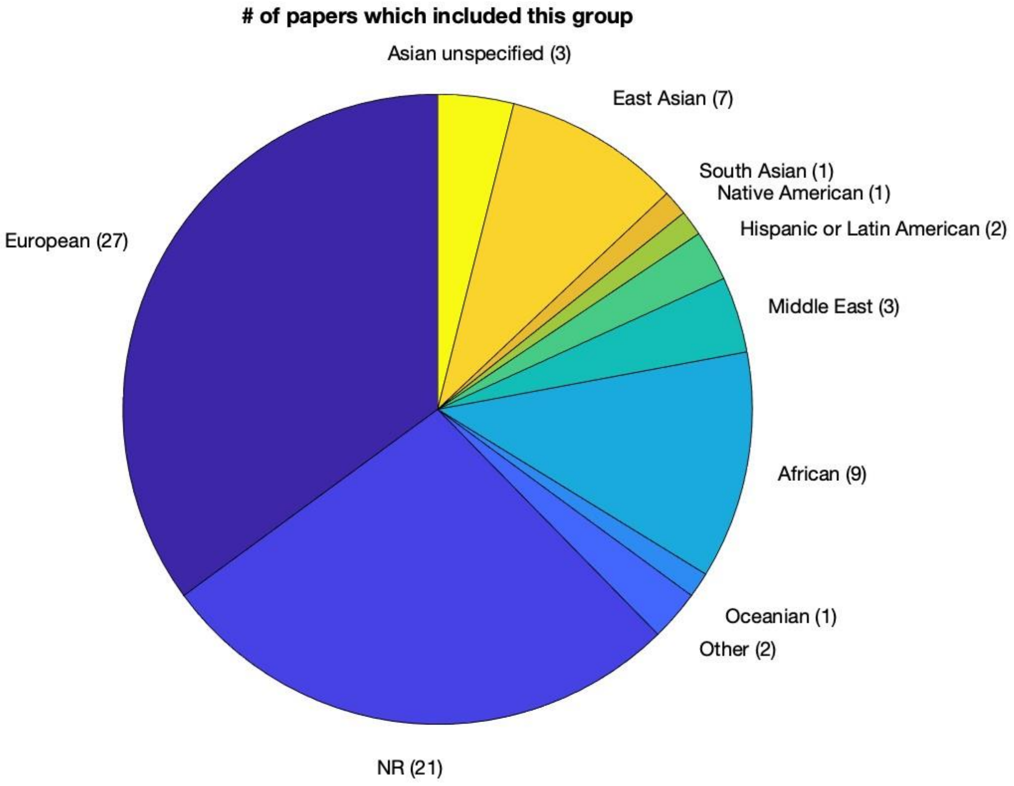

### Supplementary Figure 2

**(A)** Number of studies reported genes in GWAS associated with ASD (≥3). The 23 genes associated with ASD have been counted to be most occurring in GWAS. The green-labeled genes have been also reported to be associated with SCZ

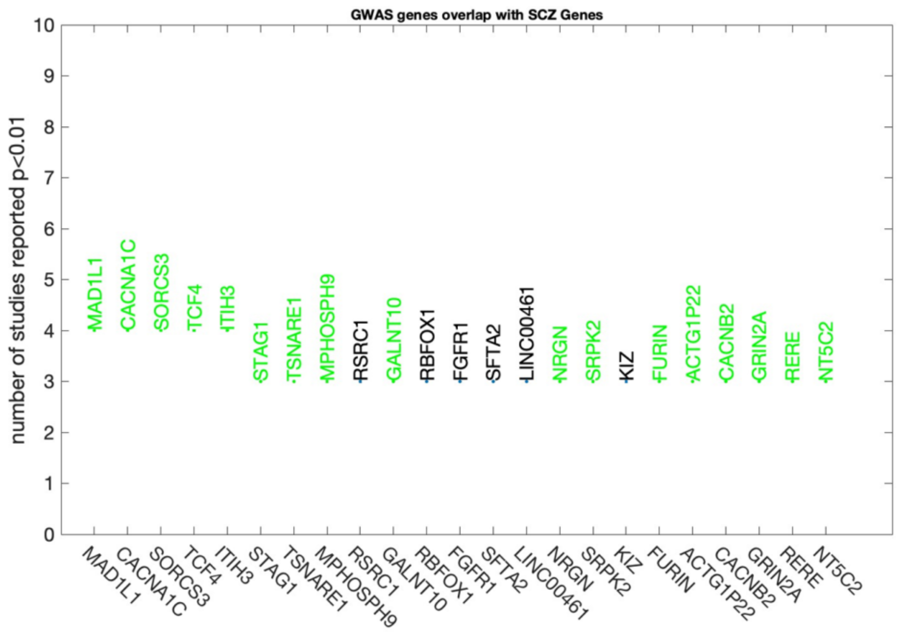


**(B)** Number of studies reported genes in GWAS associated with ASD (≥3). The red labeled genes been reported in DBD database.

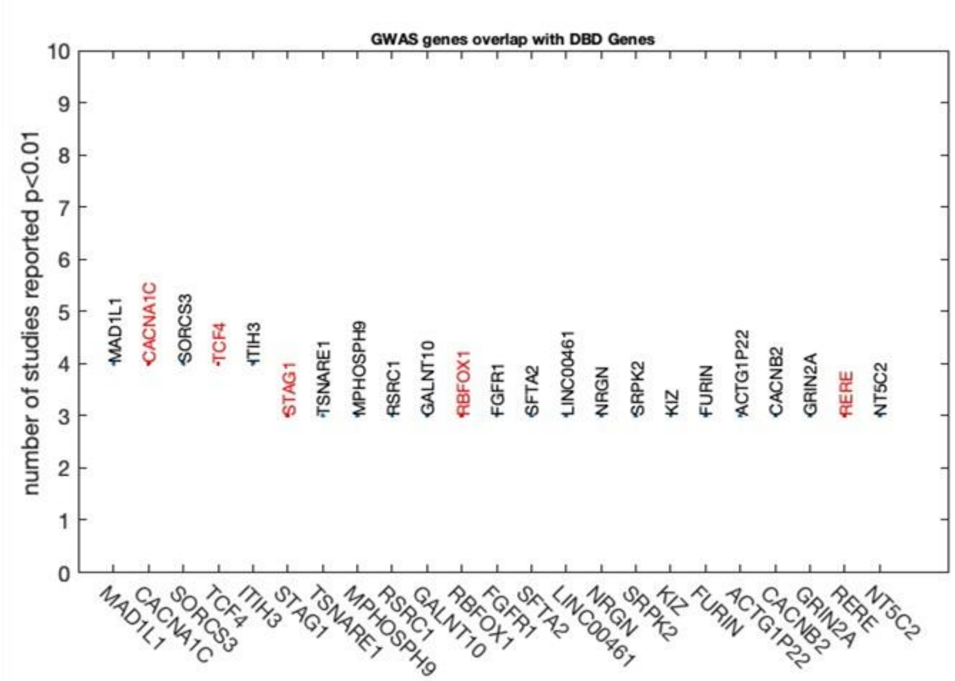

**Supplementary**
**Figure 3**

Circos plots of the distribution of SNPs in **(A)** ASD and **(B)** in shared ASD and SCZ genes. Location of each SNP associated with ASD in human chromosomes. (A) 23 most reported ASD genes with 82 SNPs in GWAS and (B) 17 shared most reported genes in ASD and SCZ with 63 SNPs in GWAS.

**A.**

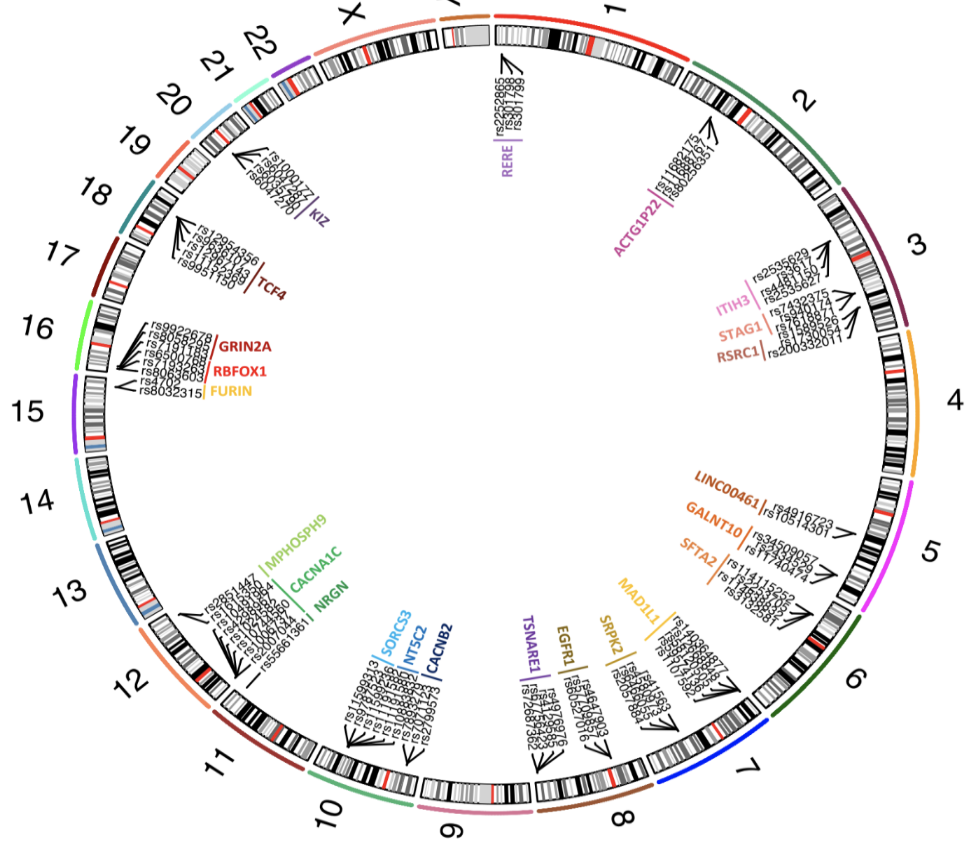


**B.**

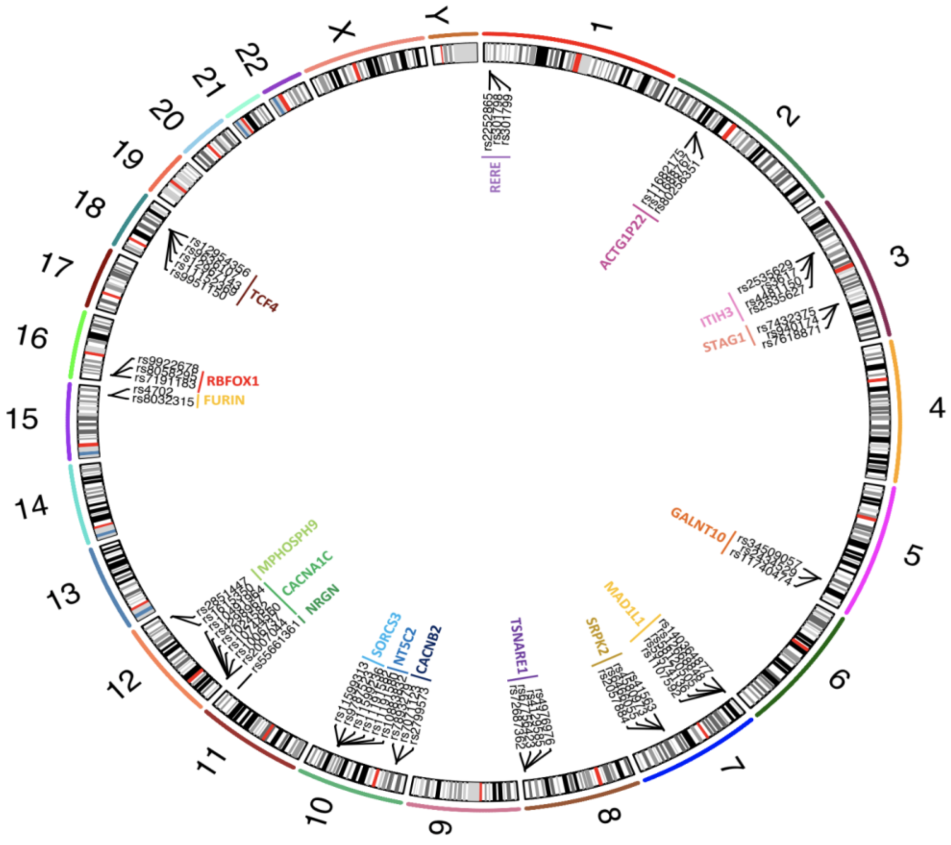
